# Strong correlation between phantomless and inline phantom-based densitometric calibration of vertebral properties from CT scans of healthy volunteers

**DOI:** 10.64898/2026.01.05.697686

**Authors:** Fiona G. Gibson, Zichu Ding, Margaret A. Paggiosi, Catherine Handforth, Janet E. Brown, Xinshan Li, Enrico Dall’Ara, Stefaan W. Verbruggen

## Abstract

Phantom calibration is currently the gold standard for calibrating CT scans and for calculating material properties of dense tissues for computational models. However, in Oncology departments and low-resource settings, it is not routine to include a calibration phantom within the scanning protocol. Therefore, retrospective scan datasets are challenging to calibrate for biomechanical investigations, precluding detailed measurements of material and mechanical properties. In this study, we compared the results from a phantomless calibration technique, where the density within each scan was independently calibrated based on known tissue densities captured within each scan (e.g. air), with those from a traditional inline phantom calibration. To do so we used scans from a cohort of healthy volunteers from the control arm of a clinical trial dataset (ANTELOPE) in which inline calibration phantoms were included. We found that, when selecting air and the aorta as regions for calibration within individual CT scans, a strong individual-specific correlation existed between bone mineral density measured in the phantomless and phantom calibrations. This indicates that the phantomless calibration method can be a useful and reliable tool for quantifying the densitometric material properties of healthy human vertebrae, and provides the opportunity for further analysis of spinal CT scans in either retrospective datasets or in low-resource clinical settings.

## 1. INTRODUCTION

To assess the patient’s risk of vertebral fracture in clinics, two imaging modalities are typically deployed, Dual Energy X-rays Absorptiometry (DXA) and Quantitative Computed Tomography (QCT). At present, DXA remains the gold standard in the clinic due to ease of use, speed, cost and the low radiation exposure (5-20 µSv by DXA vs. 60-90 µSv by QCT) (Kabayel, 2016). QCT provides images obtained from a CT scanner, where either an inline or offline (retrospective, same scanner and scanning protocol) calibration phantom could also be scanned. Calibration phantoms are used to convert the image’s Hounsfield units into equivalent bone mineral density (BMD) through a set of equations.

The role of CT technology has grown extensively due to its effectiveness in skeletal assessment for diagnosis and continuous monitoring of cancers (Duvauferrier et al., 2013). This imaging technique provides essential information for assessing spinal stability by allowing for the identification of osteopenia, lytic lesions, soft-tissue involvement and fractures (Mahnken et al., 2002). The main concern with CT use is the exposure to significantly higher doses of radiation in comparison to standard radiographs (Winterbottom and Shaw, 2009). Despite this, evidence suggests that low-dose whole body CT is effective in producing high resolution images that provide the information necessary for assessing spinal stability (Gleeson et al., 2009).

The densitometric calibration of QCT images, required to compared data across different types of scanners and protocols (Pickhardt et al., 2013; Carpenter et al., 2014), usually utilises an external phantom; however, routine QCT scans are often conducted without a densitometric calibration phantom as its usage increases the logistical burden of clinical imaging (Lee et al., 2017). To account for this, numerous methods for phantomless calibration have been developed (Michalski et al., 2020; Bartenschlager et al., 2022). One approach is to pre-calibrate the scanner using either DXA measurements or a calibration phantom and apply this general pre-calibration factor to prospective QCT scans (Budoff et al., 2013; Pickhardt et al., 2015). Even though this is an improvement from not performing any densitometric calibration, it does not consider the patient-specific differences as well as scanner and protocol changes.

The most widely used phantomless approach is to utilise internal tissues as the reference materials. The choice of tissues has been varied and depend on scan location, with some authors chose air, fat and blood (Lee et al., 2017; Schwaiger et al., 2017), while others used fat and muscle (Weaver et al., 2015; Saffarzadeh et al., 2016) and showed similar results. In particular, Bartenschlager et al. compared different combinations of two internal tissues when calibrating for vertebrae BMD, reporting the lowest error for any combination with air (<5%), particularly air and blood and the highest errors arising when using muscle in the combination (Bartenschlager et al., 2022). These authors also concluded that the use of different CT scanners did not result in significant differences in calibration outcome. However, as this study was conducted in female patients with osteoporosis, it remains to be demonstrated whether these same settings can be generalised to apply to healthy participants.

Therefore, the aim of this study was to investigate whether phantomless densitometric calibration of QCT scans can assess bone mineral density in the vertebra of healthy participants similarly to current gold standard approaches based on phantom calibration. We explored this by applying these techniques to healthy male volunteers in the control cohort of a clinical trial. Furthermore, this allowed us to compare these calibration methods across multiple time points (baseline and follow-up scans) and with two different scanners.

## 2. MATERIALS AND METHODS

### 2.1. Participant data

The dataset analysed in this study is a time series QCT dataset from the ANTELOPE clinical trial (Gibson et al., 2025). Full details of the ANTELOPE trial, which investigated the skeletal effects of androgen deprivation therapy on prostate cancer patients, and demographics of the participants have been described elsewhere (Handforth et al., 2024; Gibson et al., 2025). Ethical approval was obtained from the South Yorkshire Research Ethics Committee (IRAS ID 206171). The 25 male healthy volunteers in the trial’s control group were taken as the cohort for this study. While the trial investigated the effects of androgen deprivation therapy in prostate cancer patients (i.e. the treatment group), none of these cancer patients were included in this study. Moreover, there was no presence of vertebral fracture in any of the participants’ T12 vertebra assessed in this study. A total of 50 scans was analysed, with two scans for each of the 25 participants at baseline (0 months) and follow-up (12 months). Participant details are shown in Table 1.

**Table 1.** Participant demographics data for the 25 male healthy volunteer in the control group of the ANTELOPE clinical trial including age, height, and BMI.

| Participant ID | Age (years) | Height (cm) | BMI (kg/m<sup>2</sup>) |
| --- | --- | --- | --- |
| C01 | 74 | 192.3 | 31.4 |
| C02 | 76 | 169.0 | 22.3 |
| C03 | 77 | 180.1 | 30.1 |
| C04 | 73 | 173.3 | 26.2 |
| C05 | 78 | 170.4 | 26.9 |
| C06 | 68 | 182.8 | 27.1 |
| C08 | 53 | 184.4 | 26.9 |
| C09 | 63 | 188.1 | 29.6 |
| C10 | 70 | 161.0 | 29.3 |
| C11 | 79 | 175.0 | 30.7 |
| C12 | 78 | 179.0 | 32.1 |
| C14 | 74 | 168.2 | 26.8 |
| C15 | 80 | 175.2 | 26.3 |
| C16 | 82 | 169.0 | 29.7 |
| **C17** | 78 | 169.0 | 24.9 |
| **C19** | 75 | 180.8 | 34.9 |
| **C20** | 73 | 178.0 | 24.2 |
| **C21** | 78 | 159.8 | 22.7 |
| **C23** | 77 | 163.2 | 31.8 |
| **C24** | 73 | 184.5 | 23.9 |
| **C25** | 71 | 182.0 | 24.9 |
| **C26** | 71 | 173.0 | 20.4 |
| **C30** | 75 | 181.0 | 25.5 |
| **C31** | 71 | 171.0 | 27.6 |
| **C32** | 64 | 171.7 | 23.5 |
| **Average**<br>**(±SD)** | 73<br>$\pm 6$ | 175.9<br>$\pm 8.0$ | 27.1 $\pm 3.5$ |

### 2.2. Phantom Calibration

The QCT scan protocol for this trial included a solid inline calibration phantom (Image Analysis, Inc., Columbia, KY, USA) containing rods of 0, 0.075, and 0.15 g/cm^3^ equivalent concentration of calcium hydroxyapatite. The first cohort from 2017 (29 participants) was scanned at baseline using the GE LightSpeed VCT (GE Healthcare, Milwaukee, WI) in the radiology department at the Northern General Hospital, Sheffield, UK, whilst the follow-up scans in 2018 along with all second cohort scans (25 participants who completed all assessments) were scanned using the Toshiba Aquilion ONE (Toshiba Medical Systems, Tokyo, Japan) at the same hospital. Quality assurance was performed once per month using a Mindways phantom (Mindways Software, Inc., Austin, TX, USA) on both scanners. All scans were performed in the anteroposterior position, using the same noise index. The QCT protocol included a single scan from the cranial endplate of the T12 vertebra to the T12/L1 margin. For the GE scanner, the tube voltage was 120 kV and the mean tube current was set at 360 mA, with a voxel size of 0.937×0.937×0.625 mm^3^. For the Toshiba scanner, the tube voltage was also 120 kV, the mean tube current was set at 250 mA and a voxel size of 0.976×0.976×0.5 mm^3^.

The densitometric calibration was computed using a standard approach, which assumes a linear relationship between the average Hounsfield units (HU) and the known equivalent mean values of equivalent BMD of each rod. To do so, one region of interest (ROI) (black square boxes, Figure 1A) was defined manually within each insertion of the phantom (ImageJ) (Rasband, 1997; Schneider et al., 2012). The ROIs were defined as square regions centred within each calibration rod with length equal to half the edge length of the rod (12.5 mm). For each of the three rods, mean HU values over the same 10 slices were used to perform the linear regression analysis for calibration, Equation 1.

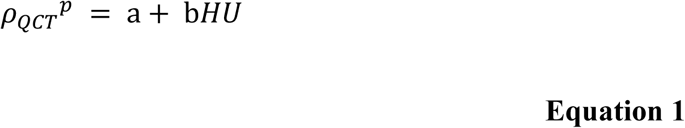

where 𝜌_𝑄𝐶𝑇_^𝑝^ represents the QCT equivalent BMD calculated using the calibration phantom, 𝐻𝑈 represents the Hounsfield unit values of the densitometric calibration law and a and b are constants from the linear regression analysis performed. This equation was applied to estimate the equivalent BMD in each voxel.

**Figure 1:**
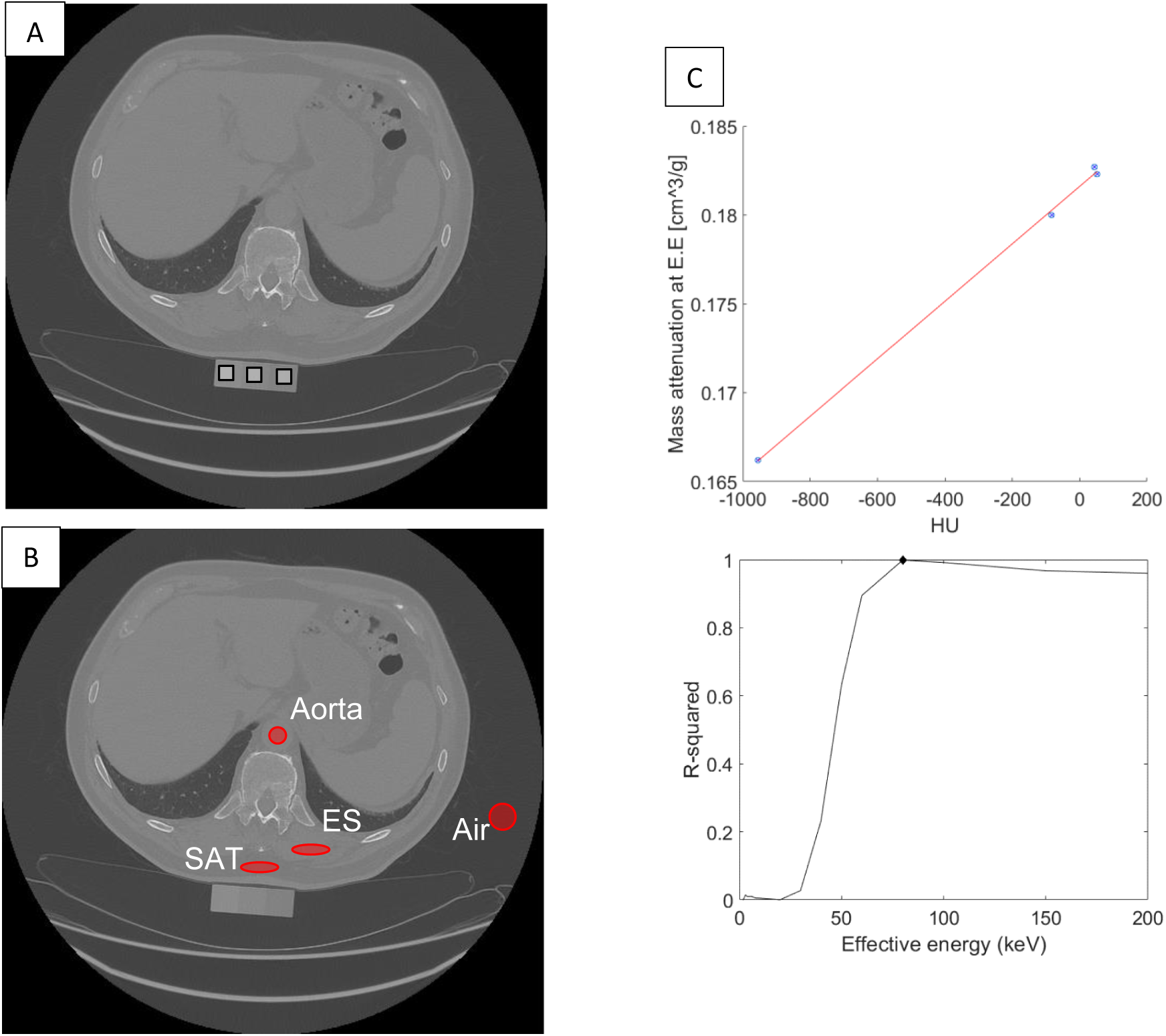
Density calibration methods for quantitative CT analysis. (A) Phantom-based calibration uses the phantom. Each calibration rod is sampled from the image to determine linear conversion between HUs and equivalent density (black square boxes). (B) In scan tissues of reference (adipose (SAT), air, blood (aorta), and skeletal muscle (ES)) are sampled adjacent to the bone of interest for internal calibration. (C) HUs and mass attenuation coefficients for each tissue (circle data points) are correlated by iterating at each effective energy (EE). Scan effective energy is determined by maximizing the coefficient of determination across all effective energies (black diamond).

### 2.3. Phantomless Calibration

To conduct the phantomless calibration, a combination of internal materials (IM) was used. From each scan, tissue ROIs for subcutaneous adipose tissue (SAT), air, aortic blood, and the erector spinae skeletal muscle (ES) were manually sampled from the scan field-of-view, as depicted in Figure 1B. To reduce influence of variations in tissue HUs across the scan field-of-view, the ROIs were placed adjacent to the bones of interest (T12 vertebra) for each tissue, and the mean HUs were determined from the tissue sample aggregated histograms of ten 2D slices, providing a compromise between the size of the ROI and size of the anatomical feature of interest. Using mass absorption coefficients (Table 2) obtained from the National Institute of Standards and Technology (www.nist.gov National Institute of Standards and Technology, NISTIR 4999), the scan effective energy was estimated by iteratively correlating the ROI-specified HUs and corresponding mass absorption coefficient at each energy level and maximizing the coefficient of determination (Millner et al., 1978), as shown in Figure 1C. This calibration for scan effective energy is necessary, as different local geometry and material properties affect the local attenuation of the X-ray beam, and this procedure allows more accurate correlation of the local HUs to the material properties (Millner et al., 1978).

**Table 2.** Mass densities of internal calibration materials obtained from the National Institute of Standards and Technology (NIST) database. ES: erector spinae muscle, SAT: subcutaneous adipose tissue.

| Material | Density (g/cm<sup>3</sup>) |
| --- | --- |
| Blood | 1.06 |
| Air | $1.205 \times 10^{-3}$ |
| ES | 1.04 |
| SAT | 0.92 |

For compounds such as hydroxyapatite (Ca10(PO4)6(OH)2) that are not tabulated in NIST, mass absorption coefficients can be calculated if the atomic mass fractions and the mass densities are known. Once the scan effective energy was determined for the scan, the mass absorption coefficients, equivalent density and measured HU values for each material were used in a two-component mass fraction model (Genant and Boyd, 1977) to calculate the associated calibration equation, Equation 4.

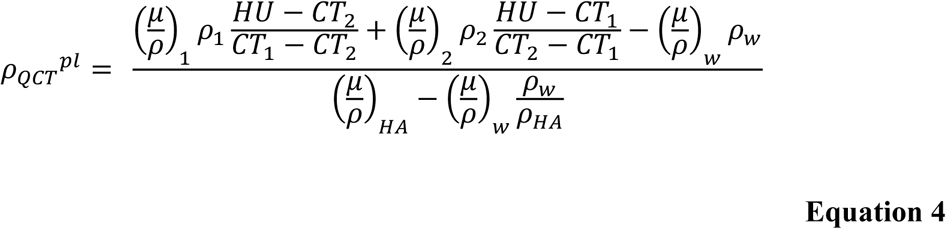

where 𝜌_𝑄𝐶𝑇_^𝑝𝑙^ represents the BMD calculated using the phantomless calibration method, 𝐻𝑈 represents the Hounsfield unit values of the densitometric calibration law, 𝐶𝑇_1_and 𝐶𝑇_2_ represent the averaged grey value of each internal material, 𝜌_1_ and 𝜌_2_ represent the density of each internal material, 𝜌_𝑤_ and 𝜌_𝐻𝐴_ represent the density of water and hydroxyapatite respectively and 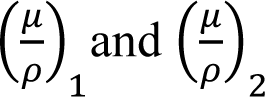 represent the mass absorption coefficient for the internal material, 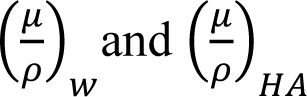 represent the mass absorption coefficients for water and hydroxyapatite respectively.

The two calculated BMD values were compared directly in order to estimate the accuracy errors for each of the 50 scans, with measured trabecular CT values of the T12 vertebra converted to BMD using both the phantom and phantomless calibration equations. Then for each scan, the difference (ΔBMDs) between the phantom based and phantomless approaches was determined.

### 2.4. Statistical Analysis

The linear regression analysis was used to compare the values of BMD calculated for each vertebra at both time points using the phantom and the phantomless calibration methods. For statistically significant regressions (p<0.05) the Spearman’s correlation coefficient (r) was reported. Steiger’s z-test was used to compare the statical significance of differences (z) between correlations established using regression analyses.

## 3. RESULTS

The systematic bias and random variability for BMD difference between phantom based and phantomless calibrations are shown in Figure 2. The largest biases were found for the combination of air and SAT (−0.0146 g/cm³, underestimation) and aorta and SAT (0.0114 g/cm³, overestimation). The air and aorta combination demonstrated the highest accuracy, with a negligible bias of 0.0006 g/cm³ and the lowest variability (SD = 0.006 g/cm³). These correspond to relative errors of 0.6% (air/aorta), 10.6% (aorta/SAT), -13.6% (air/SAT), - 10.9% (air/muscle), and -7.9% (SAT/muscle).

**Figure 2:**
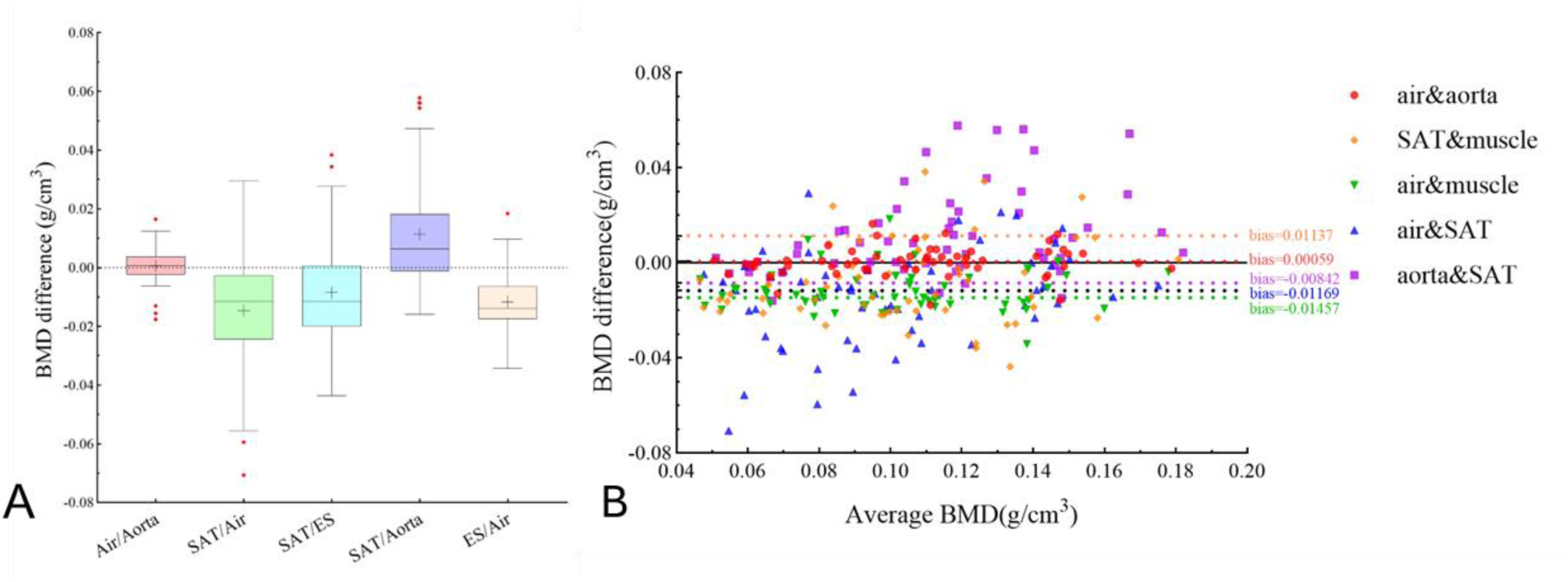
(A) BMD differences between phantomless and phantom based calibration for five different internal materials (IM) combinations. Top and bottom horizontal borders of each box indicate the 25th and 75th percentiles with their distance representing the interquartile range (IR), the black line shows the median. Red points outside the dashed lines (Whisker) are outliers with values > 1.5 × IR. (B) Test for systematic bias in the compared datasets.

Figure 3 reports the linear regressions between the phantom based and phantomless BMD values for all internal materials combinations.

**Figure 3.**
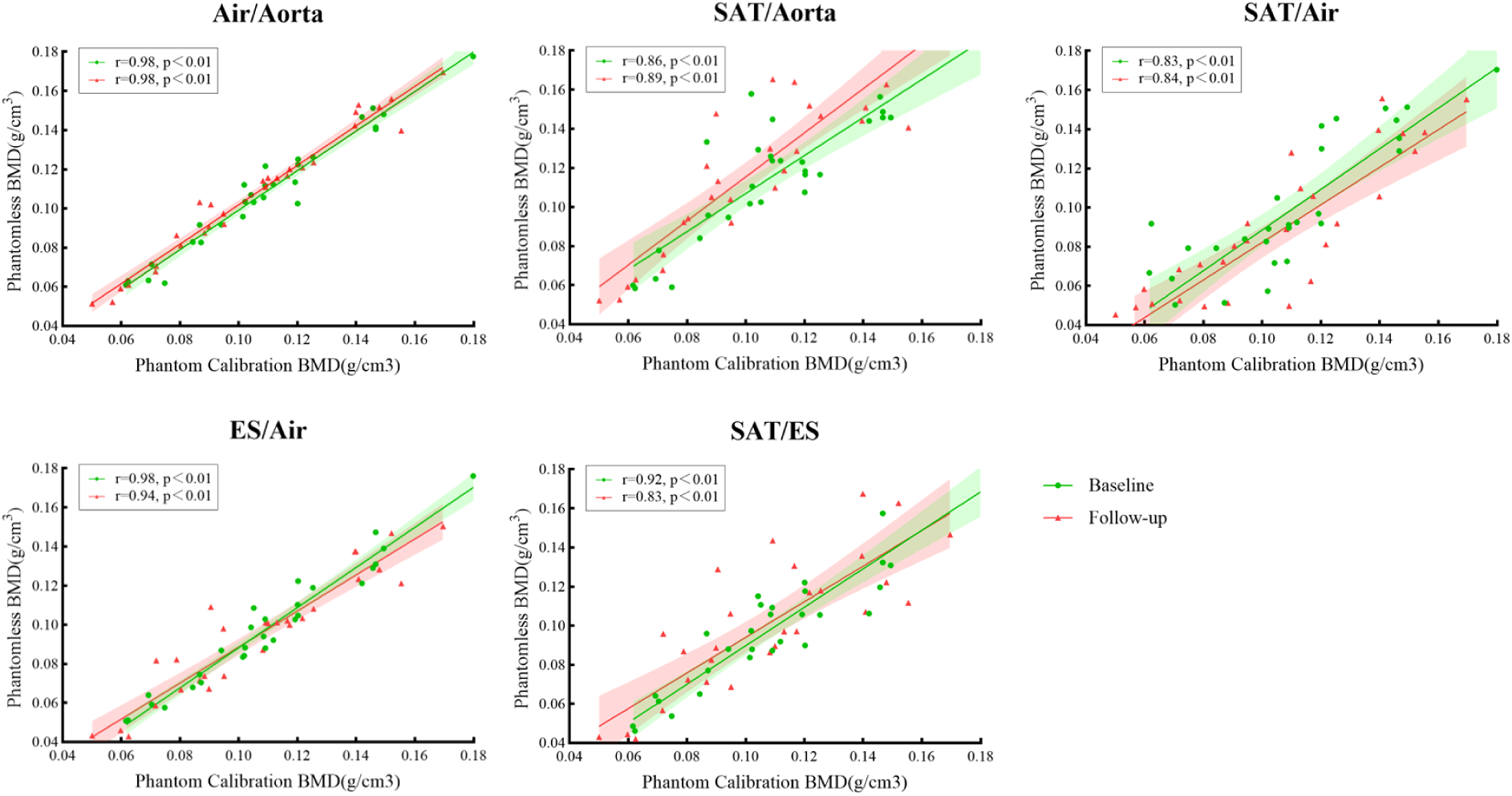
Linear regressions between the phantom and phantomless BMD for all internal material combinations, and for each group of baseline and follow-up scans

Here we found that calibration using a combination of the air in the scan, and the blood inside the aorta, provided the strongest correlation with standard phantom calibration (r=0.98, p<0.01). Additionally, using Steiger’s z-test to compare the statical significance of differences between correlations (Table 3), we found that calibration using air and aortic blood also outperformed the second-best method (air and muscle) (Steiger’s z=2.90, p<0.05). This is also inherently sensible, as the densities of air and blood are well characterised and, compared to the density of a mineralised tissue like bone, do not vary significantly from participant to participant.

**Table 3.** Comparison of regression equations between the best performer air/aorta and each of the other internal calibration materials using Steiger’s z-test.

|  | Air/Aorta vs SAT/Aorta | Air/Aorta vs SAT/Air | Air/Aorta vs ES/Air | Air/Aorta vs SAT/ES |
| --- | --- | --- | --- | --- |
| **z** | 7.5085 | 7.5512 | 2.8956 | 6.9338 |
| **p-value** | <0.0001 | <0.0001 | 0.0038 | <0.0001 |

## 4. DISCUSSION & CONCLUSION

This study compared between phantomless and phantom based calibration, and evaluated the use of different combinations of internal materials for the phantomless approach. The combination of aorta and air for calibrating the CT scans yielded the most similar volumetric BMD with a 0.41% difference when compared with the gold standard phantom method (high correlation of 0.98). These findings confirmed previous work showing that the phantomless calibration method is a useful and reliable tool for quantifying the material and mechanical properties of human vertebrae (Bartenschlager et al., 2022). Importantly, this study confirmed that these findings hold true in male healthy volunteers in clinical trial conditions, indicating that phantomless techniques may be applied more broadly, and providing the opportunity for further analysis of spinal CT scans in either retrospective datasets or in low-resource clinical settings.

A number of previous studies have demonstrated the potential of phantomless calibration for densitometric analysis of bones using QCT scans. Early work on vertebrae found that it was possible to apply phantomless calibration to the lumbar spine scans of patients injured in traffic accidents, showing excellent agreement with phantom calibration in these patients (Weaver et al., 2015; Saffarzadeh et al., 2016). Similarly, a study comparing phantomless predictions in prostate cancer patients found strong agreement in vertebral and femoral fracture risk prediction in these patients (Schwaiger et al., 2017). Further analysis by this group showed very low inter-operator variability with phantomless calibration using femur QCT scans of patients, indicating it is a reliable form of calibration (Lee et al., 2017). Most recently, Bartenschlager et al. used a retrospective QCT database of post-menopausal women to show that phantomless calibration was highly correlated with phantom based method, and that the best internal materials to use were values measured from the aorta and air (Bartenschlager et al., 2022). Our study expands upon these previous findings, showing that phantomless calibration can be applied to healthy volunteers in clinical trial conditions. We also confirmed that the most accurate calibration values were obtained when using the air and aorta as internal materials. Additionally, our study investigated changes across two different timepoints, with scans acquired using two different CT machines, demonstrating robust and broad applicability of this calibration method.

An advantage of QCT imaging of bone is that the resulting scans can be used to create 3D biomechanical models, using finite element (FE) analysis of the vertebra to estimate the bone strength. Patient-specific FE models have long been used to investigate the biomechanical response of bones to loading (Engelke et al., 2016; Schileo and Taddei, 2021). FE has been used increasingly in bones affected by diseases such as osteoporosis (Matsumoto et al., 2009) and different types of cancers including breast, colorectal and renal cell carcinoma (Costa et al., 2019). These models have also been used to study the effect of treatments and have been proven to predict vertebral strength more accurately than DXA in individuals without skeletal diseases (Crawford et al., 2003; Dall’Ara et al., 2012) and with osteoporosis (Imai, 2015). We have recently applied such FE modelling using the ANTELOPE trial dataset reported here, using phantom based calibration to investigate the effects of androgen deprivation therapy on vertebral biomechanics (Gibson et al., 2025). In that study we found that simple BMD changes due to treatment did not capture the full extent of the degradation in vertebral strength evidenced in the FE models. Therefore it remains to be seen whether the similar correlation between phantom and phantomless calibration shown here, and elsewhere (Bartenschlager et al., 2022), will propagate through into resulting FE predictions of vertebral biomechanics.

## ACKNOWLEDGMENTS

We would like to thank all the ANTELOPE trial participants, and the Weston Park Cancer charity for funding the ANTELOPE trial.

## Author contributions

FG - Data curation, Formal analysis, Investigation, Methodology, Validation, Visualisation, Writing-original draft, Writing – review and editing

ZD - Data curation, Formal analysis, Visualisation, Writing-original draft, Writing – review and editing

CH – Data curation, Formal analysis, Investigation, Methodology, Validation, Visualisation, MAP – Data curation, Formal analysis, Investigation, Methodology, Project administration, Resources, Supervision, Validation, Writing – review and editing

JEB – Conceptualisation, Data curation, Funding acquisition, Investigation, Methodology, Project administration, Resources, Supervision, Validation, Visualisation, Writing-original draft, Writing – review and editing

XL – Methodology, Project administration, Resources, Supervision, Validation, Visualisation, Writing – review and editing

ED – Conceptualisation, Methodology, Project administration, Resources, Supervision, Validation, Visualisation, Writing – review and editing

SV - Conceptualisation, Funding acquisition, Investigation, Methodology, Project administration, Resources, Supervision, Validation, Visualisation, Writing-original draft, Writing – review and editing

## Data availability

The data that support the findings of this study are available on reasonable request from the corresponding author. All requests will be reviewed by relevant stakeholders on the basis of a controlled access approach. The data are not publicly available due to ethical restrictions.

